# Respiratory disease and virus shedding in rhesus macaques inoculated with SARS-CoV-2

**DOI:** 10.1101/2020.03.21.001628

**Authors:** Vincent J. Munster, Friederike Feldmann, Brandi N. Williamson, Neeltje van Doremalen, Lizzette Pérez-Pérez, Jonathan Schulz, Kimberly Meade-White, Atsushi Okumura, Julie Callison, Beniah Brumbaugh, Victoria A. Avanzato, Rebecca Rosenke, Patrick W. Hanley, Greg Saturday, Dana Scott, Elizabeth R. Fischer, Emmie de Wit

**Affiliations:** Laboratory of Virology, National Institutes of Health, Hamilton, MT, United States of America; Rocky Mountain Veterinary Branch, National Institutes of Health, Hamilton, MT, United States of America; Research Technologies Branch, National Institute of Allergy and Infectious Diseases, National Institutes of Health, Hamilton, MT, United States of America

## Abstract

An outbreak of a novel coronavirus, now named SARS-CoV-2, causing respiratory disease and a ∼2% case fatality rate started in Wuhan, China in December 2019. Following unprecedented rapid global spread, the World Health Organization declared COVID-19 a pandemic on March 11, 2020. Although data on disease in humans are emerging at a steady pace, certain aspects of the pathogenesis of SARS-CoV-2 can only be studied in detail in animal models, where repeated sampling and tissue collection is possible. Here, we show that SARS-CoV-2 causes respiratory disease in infected rhesus macaques, with disease lasting 8-16 days. Pulmonary infiltrates, a hallmark of human disease, were visible in lung radiographs of all animals. High viral loads were detected in swabs from the nose and throat of all animals as well as in bronchoalveolar lavages; in one animal we observed prolonged rectal shedding. Taken together, the rhesus macaque recapitulates moderate disease observed in the majority of human cases. The establishment of the rhesus macaque as a model of COVID-19 will increase our understanding of the pathogenesis of this disease and will aid development and testing of medical countermeasures.

## Main

A novel coronavirus, designated SARS-CoV-2, emerged in Wuhan, China at the end of 2019^1,2^ and quickly spread across the globe. The World Health Organization declared a public health emergency of international concern on January 30, 2020 and, when the spread of SARS-CoV-2 continued at a rapid pace, a pandemic on March 11, 2020. As of March 19, 2020, more than 240,000 cases have been identified in at least 160 countries, including more than 10,000 fatal cases^3^. Coronavirus Disease 2019 (COVID-19) has a broad clinical spectrum ranging from mild to severe cases^4-6^. Patients with COVID-pneumonia presented mainly with fever, fatigue, dyspnea and cough^7-9^. Rapidly progressing pneumonia, with bilateral opacities on x-ray or patchy shadows and ground glass opacities by CT scan were observed in the lungs of patients with COVID-19 pneumonia^2,6,10^. Older patients with comorbidities are at highest risk for adverse outcome of COVID-19^5,7^. SARS-CoV-2 has been detected in upper and lower respiratory tract samples from patients, with high viral loads in upper respiratory tract samples^11-13^. In addition, virus has been detected in feces and blood but not in urine^13,14^.

Non-human primate models that recapitulate aspects of human disease are essential for our understanding of the pathogenic processes involved in severe respiratory disease. In addition, these models are crucial for the development of medical countermeasures such as vaccines and antivirals. Previously, we successfully established rhesus macaques as an animal model for the pre-clinical development of vaccines and antivirals against the related Middle East Respiratory Syndrome Coronavirus (MERS-CoV)^15-17^. Here, we provide a detailed description of a rhesus macaque model of moderate COVID-19.

### Clinical, respiratory disease upon inoculation with SARS-CoV-2

Eight adult rhesus macaques were inoculated with a total dose of 2.6×10^6^ TCID50 of SARS-CoV-2 isolate nCoV-WA1-2020^18^ via a combination of intratracheal, intranasal, ocular and oral routes. On day 1 post inoculation (dpi), all animals showed changes in respiratory pattern and piloerection, as reflected in their clinical scores (Fig. 1a). Other observed signs of disease included reduced appetite, hunched posture, pale appearance and dehydration (Table S1). Disease signs persisted for more than a week, with all animals completely recovered between 9 and 17 dpi (Fig. 1a and Table S1). Weight loss was observed in all animals (Fig. 1b); body temperatures spiked on 1 dpi but returned to normal levels thereafter (Fig. 1c). Under anesthesia, the animals did not show increased respiration; however, all animals showed irregular respiration patterns (Fig. 1d). Radiographs showed pulmonary infiltrates in all animals starting on 1 dpi with mild pulmonary infiltration primarily in the lower lung lobes. By 3 dpi, progression of mild pulmonary infiltration was noted into other lung lobes although still primarily in the caudal lung lobes (Fig. 1e). In one animal, pulmonary infiltrates were observed from 1-12 dpi (Fig. 2). This animal had a moderate pulmonary infiltration in the upper right lung lobe starting on 3 dpi, with resolution by 7 dpi. However, on 10 dpi, pulmonary infiltration appeared to reappear in previously affected lung lobes. These signs began to resolve by 14 dpi (Fig. 2).

**Figure 1.**
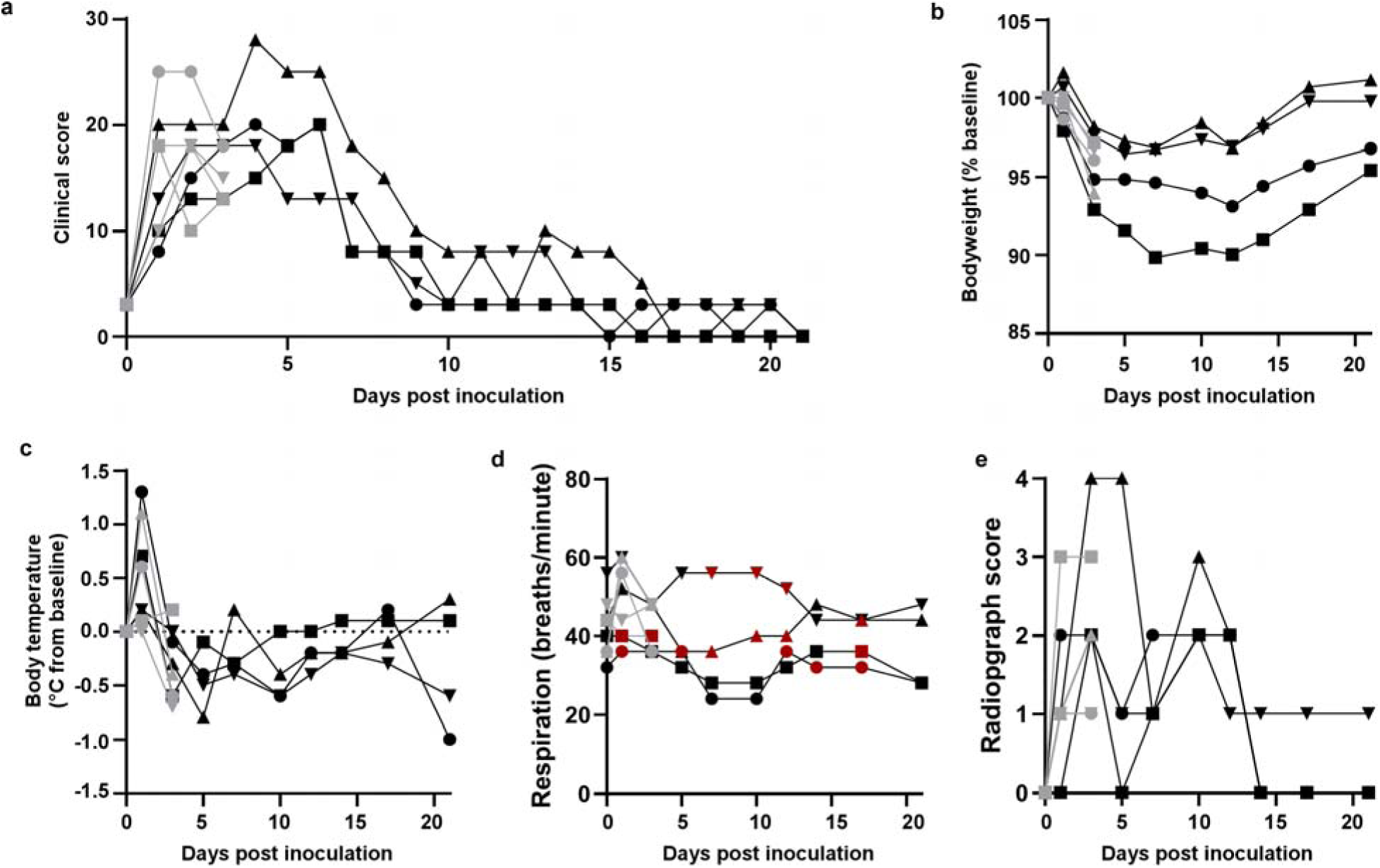
Rhesus macaques infected with SARS-CoV-2 develop respiratory disease. Eight adult rhesus macaques were inoculated with SARS-CoV-2 isolate nCoV-WA1-2020. After inoculation, animals were observed for disease signs and scored according to a pre-established clinical scoring sheet (a). On days when clinical exams were performed, body weight (b), and body temperature (c) were measured. Respiration rate was measured, and breathing pattern was recorded, with irregular respiration patterns indicated in red (d). Ventro-dorsal and lateral radiographs were taken on clinical exam days and scored for the presence of pulmonary infiltrates by a clinical veterinarian according to a standard scoring system (0: normal; 1: mild interstitial pulmonary infiltrates; 2: moderate pulmonary infiltrates perhaps with partial cardiac border effacement and small areas of pulmonary consolidation; 3: severe interstitial infiltrates, large areas of pulmonary consolidation, alveolar patterns and air bronchograms). Individual lobes were scored and scores per animal per day were totaled and displayed (e). Grey: data derived from animals euthanized on 3 dpi; black: data derived from animals euthanized on 21 dpi. Identical symbols have been used to denote identical animals throughout the figures in this manuscript.

**Figure 2.**
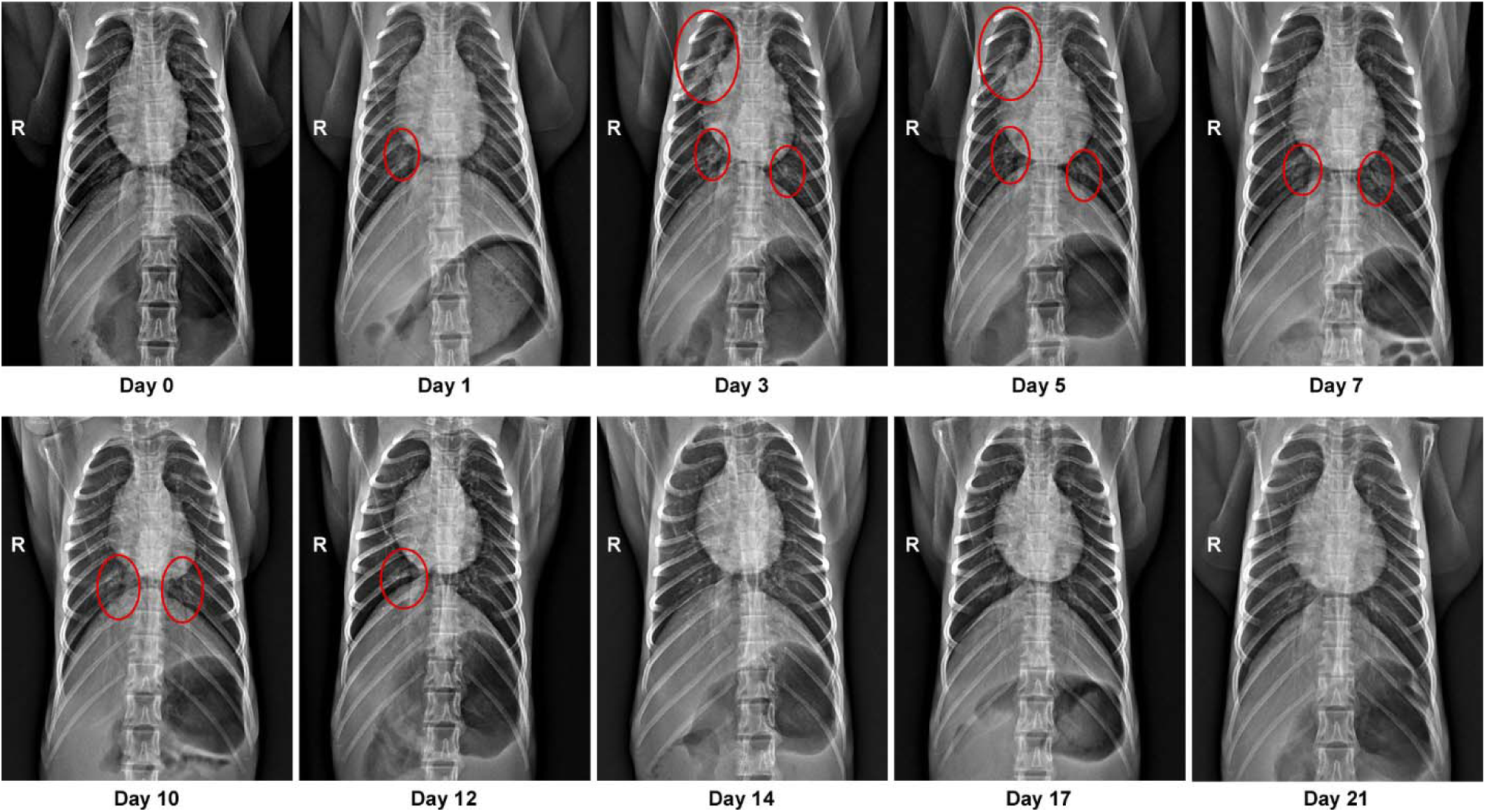
Pulmonary infiltrates in a rhesus macaque after inoculation. Radiographs show the progression of pulmonary infiltrates throughout the study in a single animal. Of note, this animal is denoted with a black triangle throughout the manuscript. Circles indicate areas of mild to moderate pulmonary infiltrates. A marker ^®^ indicates right side of the animal.

Hematologic analysis of blood collected during clinical exams showed evidence of a stress leukogram^19^ by 1 dpi in the majority of animals characterized by leukocytosis, neutrophilia, monocytosis and lymphopenia (Fig. S1). Lymphocytes and monocytes returned to baseline after 1 dpi. Neutrophils decreased in all animals by 3 dpi and continued to decline through 5 dpi; neutropenia was observed in 2 of 4 animals. Neutrophils continued to be depressed through 10 dpi and began to recover thereafter. On 1 dpi, decreased hematocrit, red blood cell counts and hemoglobin were observed in all animals (Fig. S1). In 3 of 4 animals, the values remained low or continued to decrease until 5 dpi. In addition, reticulocyte percentages and counts also decreased during this time period. At 5 dpi, two of the four animals had a normocytic, normochromic non-regenerative anemia consistent with anemia of critical illness. Although the anemia appeared to stabilize and parameters improved slightly, both animals did not return to their original baselines by 21 dpi. Blood chemistry analysis revealed no values outside normal range throughout the experiment (Table S2).

Serum was analyzed for changes in cytokine and chemokine levels at different time points after inoculation. Statistically significant changes were only observed on 1 dpi, with increases in IL1ra, IL6, IL10, IL15, MCP-1, MIP-1b, and on 3 dpi a small but statistically significant decrease in TGFα was observed (Fig. S2). Although changes occurred in the levels of some of these cytokines later after inoculation, these mostly occurred in single animals and were thus not statistically significant (Fig. S2).

### High viral loads in respiratory samples

During clinical exams, nose, throat, rectal and urogenital swabs were collected (Fig. 3a). Virus shedding was highest from the nose; virus could be isolated from swabs collected on 1 and 3 dpi, but not thereafter. Viral loads were high in throat swabs immediately after inoculation but were less consistent than nose swabs thereafter; in one animal throat swabs were positive on 1 and 10 dpi but not on any of the sampling dates in between. One animal showed prolonged shedding of viral RNA in rectal swabs from 7-17 dpi; infectious virus could not be isolated from these swabs (Fig. 3a). Urogenital swabs remained negative in all animals throughout the study. On 1, 3 and 5 dpi bronchoalveolar lavages (BAL) were performed on the 4 animals in the group euthanized on 21 dpi as a measure of virus replication in the lower respiratory tract. High viral loads were detected in BAL fluid in all animals on all three time points; infectious virus could only be isolated in BAL fluid collected on 1 and 3 dpi. No viral RNA could be detected in blood throughout the study (Fig. 3c) or urine collected at 3 and 21 dpi (Fig. 2d).

**Figure 3.**
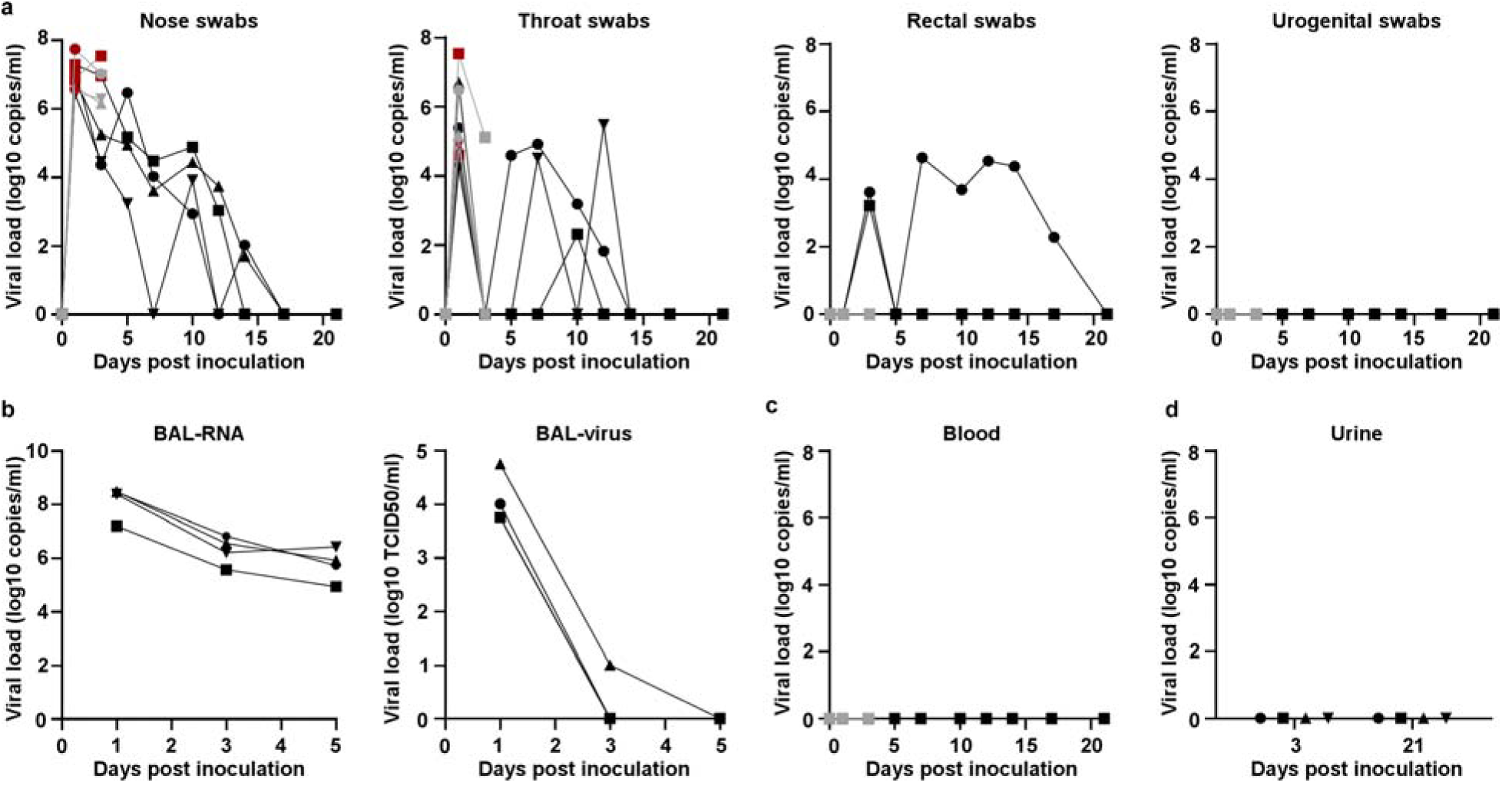
Viral loads in respiratory samples and bodily fluids. Eight adult rhesus macaques were inoculated with SARS-CoV-2 isolate nCoV-WA1-2020. After inoculation, clinical exams were performed during which nose, throat, rectal and urogenital swabs were collected; viral loads in these samples were determined by qRT-PCR (a). On 1, 3, and 5 dpi, bronchoalveolar lavages were performed on the 4 animals remaining in the study through 21 dpi; viral loads and virus titers were determined in these samples. Viral loads were also determined in blood collected during clinical exams (c) and in urine collected at necropsy on 3 and 21 (d). Grey: data derived from animals euthanized on 3 dpi; black: data derived from animals euthanized on 21 dpi; red: virus was isolated from these samples. Identical symbols have been used to denote identical animals throughout the figures in this manuscript.

### Interstitial pneumonia centered on terminal bronchioles

On 3 and 21 dpi, one group of 4 animals was euthanized and necropsies were performed. On 3 dpi, varying degrees of gross lung lesions were observed in all animals (Fig. 4a and c). By 21 dpi, gross lesions were still visible in the lungs of 2 of 4 animals (Fig. 4b and c). Additionally, all animals had an increased lung weight:body weight ratio (Fig. 4d), indicative of pulmonary edema. Histologically, 3 of the 4 animals euthanized on 3 dpi developed some degree of pulmonary pathology. Lesions were multifocal (Fig. S3), mild to moderate, interstitial pneumonia that frequently centered on terminal bronchioles. The pneumonia was characterized by thickening of alveolar septae by edema fluid and fibrin and small to moderate numbers of macrophages and fewer neutrophils. Alveoli contained small numbers of pulmonary macrophages and neutrophils. Lungs with moderate changes also had alveolar edema and fibrin with formation of hyaline membranes. There was minimal type II pneumocyte hyperplasia. Occasionally, bronchioles showed necrosis, loss and attenuation of the epithelium with infiltrates of neutrophils, macrophages and eosinophils. Multifocally, there were perivascular infiltrates of small numbers of lymphocytes forming perivascular cuffs. Three of 4 animals on 3 dpi had fibrous adhesions of the lung to the pleura. Histologic evaluation shows these to be composed of mature collagen interspersed with small blood vessels; therefore, this is most likely a chronic change rather than related to SARS-CoV-2 infection.

**Figure 4.**
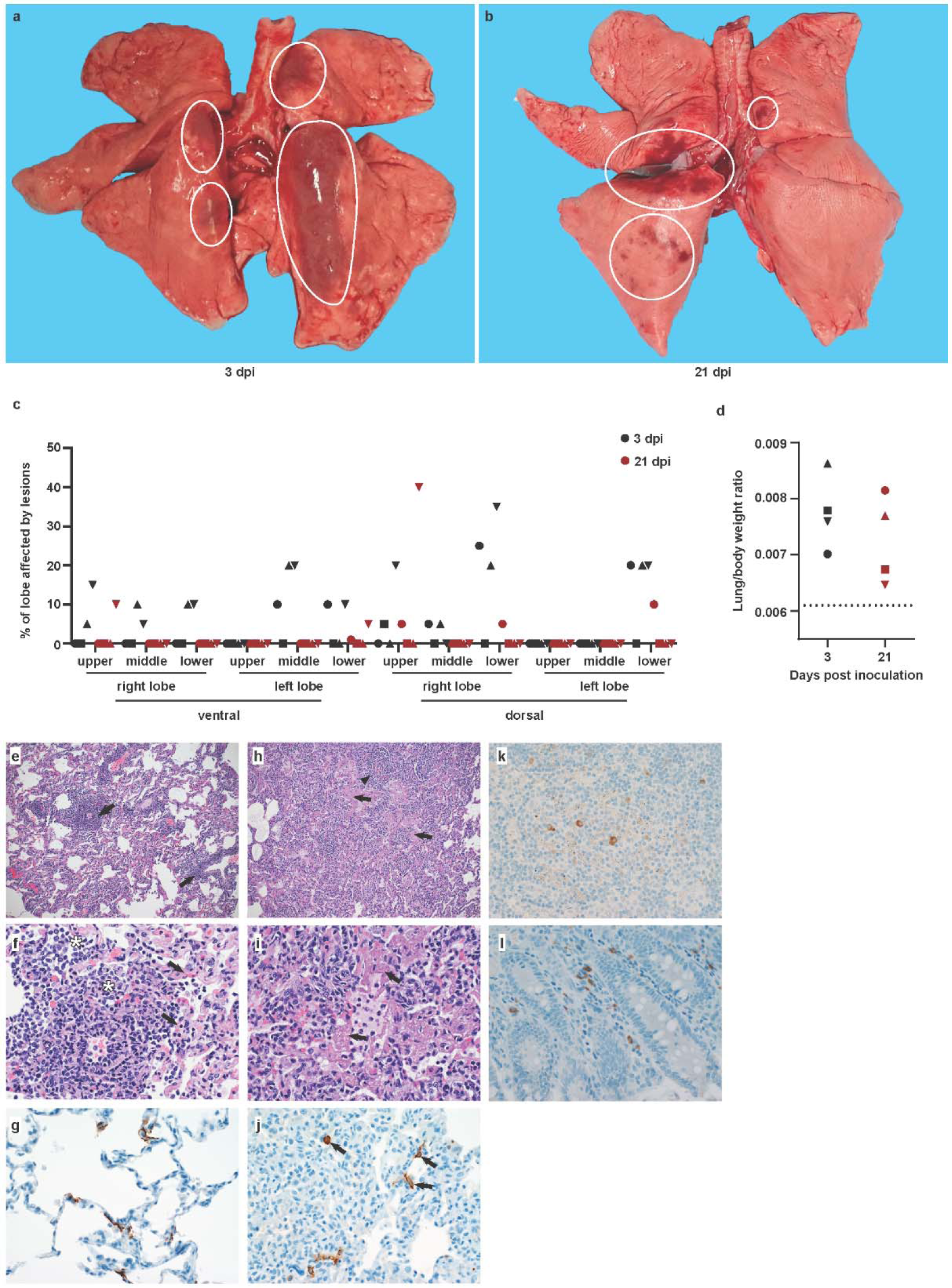
Pathological changes in rhesus macaques infected with SARS-CoV-2. Eight adult rhesus macaques were inoculated with SARS-CoV-2 isolate nCoV-WA1-2020. Four animals were euthanized on 3 dpi, and 4 animals on 21 dpi. Upon gross examination, lungs showed focal areas of hilar consolidation and hyperemia (circles) on 3 dpi (a) and multifocal, random consolidation and hyperemia (circles) on 21 dpi (b). At necropsy, the percentage of the area of the lungs affected by gross lesions was estimated by a board-certified veterinary pathologist (c), and lung weight to bodyweight ratio was calculated as a measure for pulmonary edema. (d). The dotted line represents baseline ratio calculated from an in-house collection of rhesus macaque lung and bodyweights from animals with grossly normal lungs. Histological analysis was performed on tissues collected at 3 dpi (e-i). (e) Pulmonary vessels are surrounded by moderate numbers of inflammatory cells (arrows) Magnification 100x. (f) Alveoli are filled with small to moderate numbers of inflammatory cells (asterisks). Adjacent alveolar interstitium (arrows) is thickened by edema, fibrin and inflammatory cells. Magnification 400x. (g) SARS-CoV-2 antigen is detected by immunohistochemistry in type I pneumocytes. Magnification 400x. (h) Pulmonary vessels are bounded by inflammatory cells (arrowhead) and hyaline membranes (arrows) line alveolar spaces. Magnification 100x. (i) Hyaline membranes line alveoli (arrows). Magnification 400x. (j) SARS-CoV-2 antigen is detected by immunohistochemistry in type I pneumocytes (asterisk) and type II pneumocytes (arrow) as well as alveolar macrophages (arrowheads). Magnification 400x. (k) SARS-CoV-2 antigen is detected by immunohistochemistry in mediastinal lymph node. Magnification 400x. (l) SARS-CoV-2 antigen is detected by immunohistochemistry in macrophages and lymphocytes in the lamina propria of the cecum. Magnification 400x.

Immunohistochemistry using a mAb against SARS-CoV demonstrated viral antigen in small numbers of type I and II pneumocytes, as well as alveolar macrophages. Antigen-positive macrophages were detected in mediastinal lymph nodes of 3 of 4 animals. Interestingly, small numbers of antigen-positive lymphocytes and macrophages were also detected in the lamina propria of the intestinal tract of all 4 animals. In one animal, all collected tissues of the gastrointestinal tract showed these antigen-positive mononuclear cells (Fig. S4). Ultrastructural analysis of lung tissue by transmission electron microscopy confirmed the histologic diagnosis of interstitial pneumonia. The alveolar interstitial space was greatly expanded by edema, fibrin, macrophages and neutrophils (Fig. 5a). The subepithelial basement membrane was unaffected and maintained a consistent thickness and electron density. Occasionally, type I pneumocytes are separated from the basement membrane by edema; the resulting space may contain virions. Affected type I pneumocyte are lined by small to moderate numbers of virions 90-160 nm in diameter with an electron dense core bound by a less dense capsid (Fig5b-e). Alveolar spaces adjacent to affected pneumocytes are filled with a granular, moderately electron dense material that is consistent with edema fluid.

**Figure 5.**
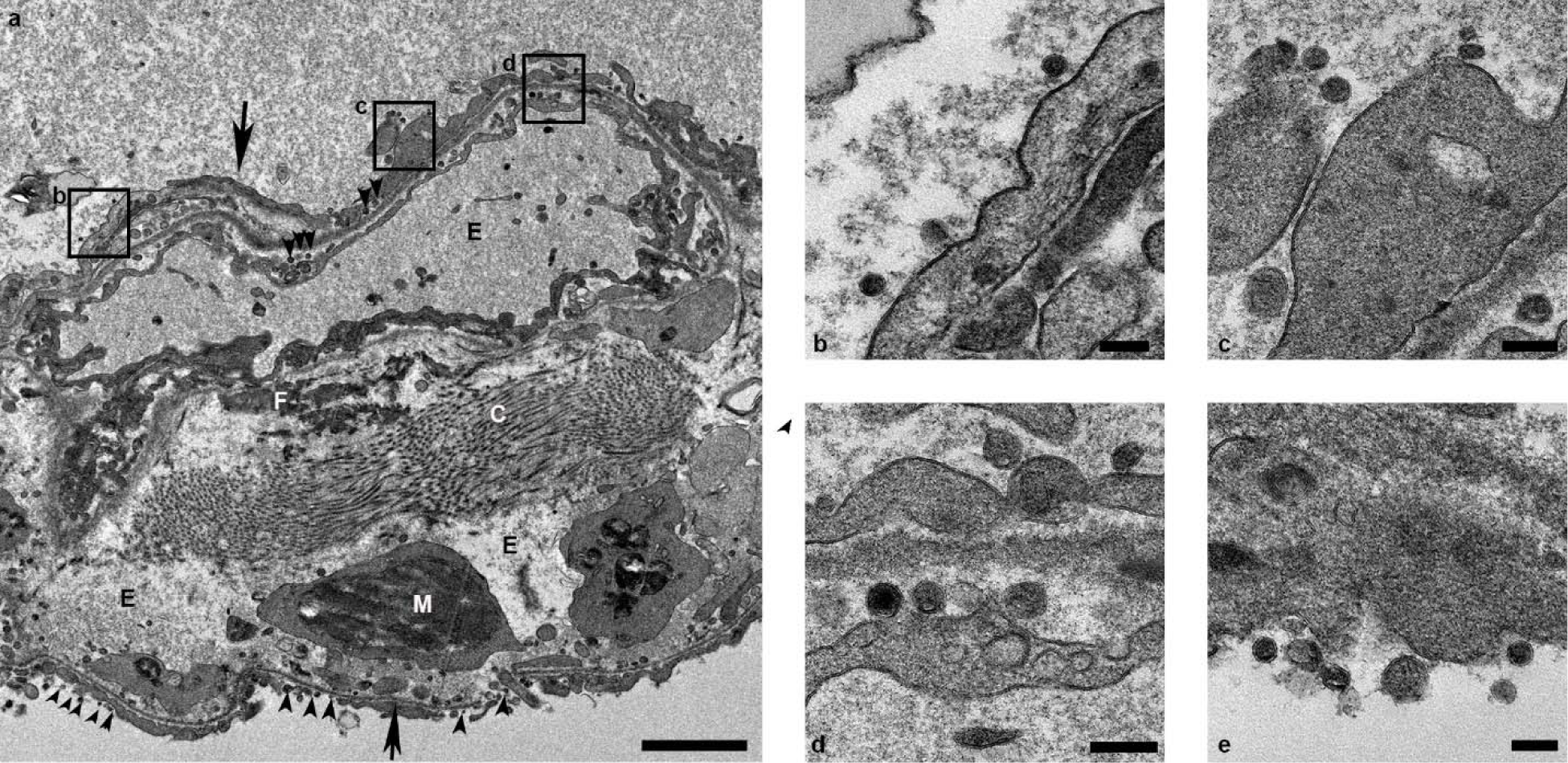
Ultrastructural analysis of lungs of rhesus macaques infected with SARS-CoV-2. Lung tissue collected on 3 dpi was analyzed by transmission electron microscopy. The alveolar interstitium is expanded by edema (E), fibrin (F) and mononuclear (M) inflammatory cells (a). Normal collagen fibers (c) and multiple virions (arrowheads) line type I pneumocytes (arrows). Boxes in (a) indicate areas enlarged in (b-d). Scale bar in (a) represents 2µm, scale bars in (b-e) represent 0.2 µm.

### SARS-CoV-2 mainly replicates in the respiratory tract

All tissues (n=37) collected at necropsy were analyzed for the presence of viral RNA. On 3 dpi, high viral loads were detected in the lungs of all animals (Fig. 6a); virus could be isolated from the lungs of all 4 animals at this time. Additionally, viral RNA could be detected in other samples throughout the respiratory tract (Fig. 6), as well as in lymphoid tissue and in gastrointestinal tissues from several animals. Viral RNA could not be detected in major organs including the central nervous system. In an attempt to distinguish viral RNA derived from respiratory secretions from active virus replication, all samples with presence of viral RNA were also tested for the presence of viral mRNA (Fig. 6). Viral mRNA was detected in all respiratory tissues could not be detected in any but one of the gastrointestinal tissues, indicating that virus replication in these tissues seems unlikely, although we can’t exclude it due to limited size of the tested samples. By 21 dpi, viral RNA, but not mRNA, could still be detected in tissues from all 4 animals (Fig. S5). In the animal with prolonged rectal shedding (Fig. 3a) and enlarged mesenteric lymph nodes on 21 dpi (Table S1), viral RNA could be detected in the mesenteric lymph nodes but not in any gastrointestinal tissues.

**Figure 6.**
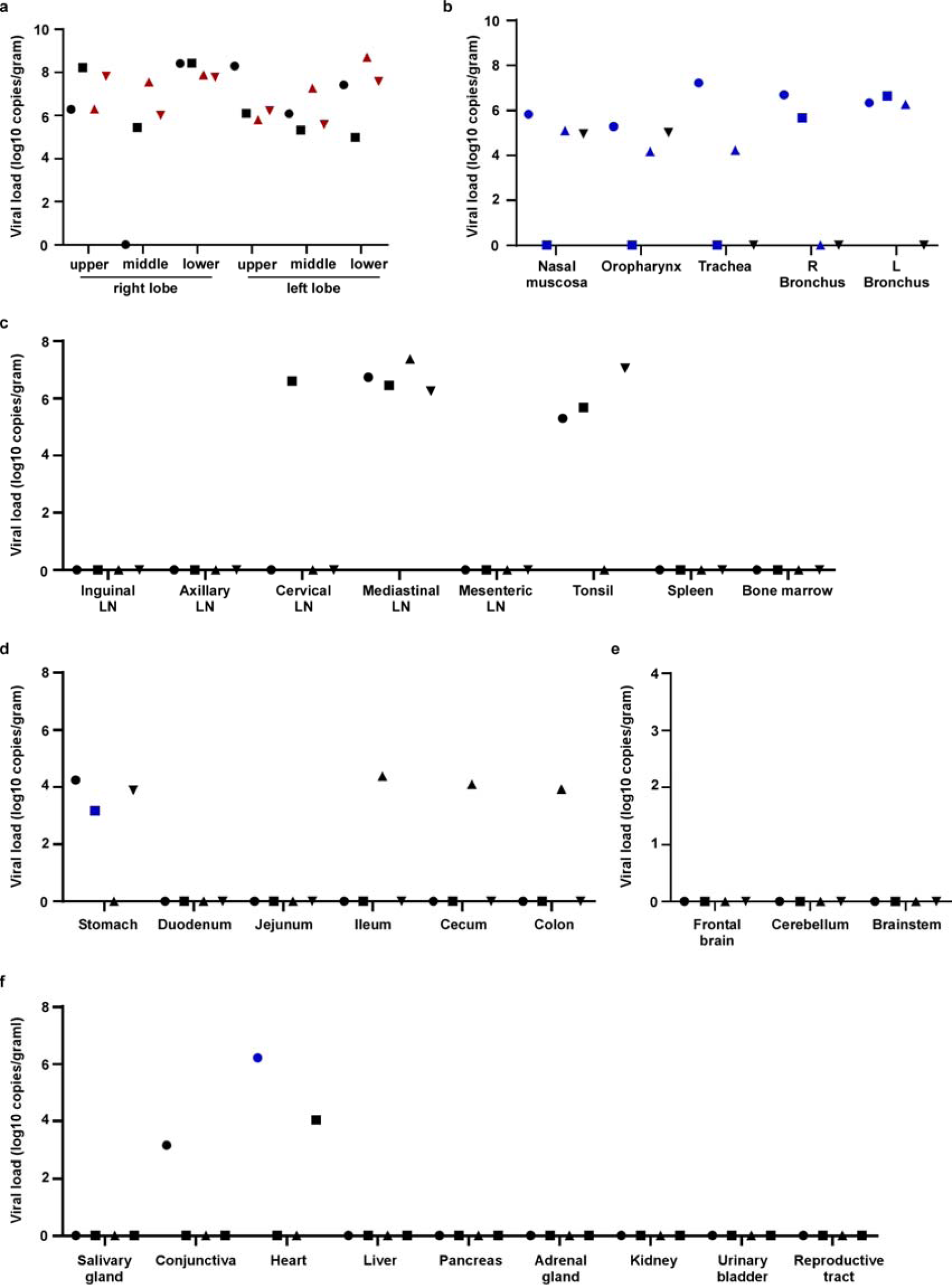
Viral loads in tissues collected from rhesus macaques infected with SARS-CoV-2. Four adult rhesus macaques were inoculated with SARS-CoV-2 isolate nCoV-WA1-2020 and euthanized on 3 dpi. Thirty-seven tissues were collected at necropsy and analyzed for the presence of viral RNA by qRT-PCR. Tissues are grouped by lung lobes (a), with red symbols indicating tissues from which virus could be isolated in Vero E6 cells; other tissues from the respiratory tract (b); lymphoid tissues (c); gastrointestinal tissues (d); the central nervous system (e) and remaining tissues (f). Blue symbols in b-e indicate that viral mRNA was also detected in these tissues. Identical symbols have been used to denote identical animals throughout the figures in this manuscript. R: right; L: left; LN: lymph node.

### Serology

Serum was analyzed for the development of IgG against SARS-CoV spike in ELISA. By 10 dpi, all four animals had seroconverted to SARS-CoV-2 spike. At 21 dpi ELISA titers ranged between 1600 – 3200 for all four animals. Neutralizing responses also started to appear at 10 dpi, and ranged from 10 – 60 at 21 dpi (Figure S6). Interestingly, the animal with the lowest and latest neutralizing antibody response was the animal with prolonged viral shedding from the intestinal tract.

## Discussion

After SARS-CoV and MERS-CoV, SARS-CoV-2 is the third coronavirus capable of causing severe respiratory disease in the human population to emerge in the past 17 years^2,20^. In contrast to the emergence of SARS-CoV and MERS-CoV, clinical data have become available in real-time. COVID-19 has a broad spectrum of clinical manifestations, ranging from asymptomatic to mild to severe^5,6,8,9,14,21^. Patients present with influenza-like symptoms such as a fever and shortness of breath and may subsequently develop pneumonia requiring mechanical ventilation and support in an intensive care unit^9^. Also similar to SARS-CoV-1 and MERS-CoV, comorbidities such as hypertension and diabetes appear to play an important role in adverse outcome of COVID-19^8,22,23^. Advanced age and chronic conditions in particular are indicators of a negative outcome^5,7-9,21^. A recent analysis of 1099 COVID-19 cases from China showed that approximately 5% of diagnosed patients developed severe pneumonia requiring ICU attendance, 2.3% required mechanical ventilation and 1.4% died^9^. The transient, moderate disease developed within our rhesus macaque model is thus in line with the vast majority of human COVID-19 cases. Pulmonary infiltrates on radiographs, a hallmark of human infection^2,4,6,7,9,10,21^, were observed in all macaques. The shedding pattern observed in rhesus macaques is strikingly similar to that observed in humans^11,12^. In humans, consistent high SARS-CoV-2 shedding was observed from the upper and lower respiratory tract, frequent intermediate shedding from the intestinal tract and sporadic detection in blood^13^. In rhesus macaques, high viral loads were detected in nasal swabs that remained positive for up to 14 days after inoculation; throat swabs were intermittently positive for up to 12 days and rectal swabs up to 17 days. Similar to what has been observed in humans, shedding of SARS-CoV-2 continued after resolution of clinical symptoms and radiologic abnormalities ^24^. Limited histopathology is available from COVID-19 patients^25,26^. Our analysis of the histopathological changes observed in rhesus macaques, suggests that they resemble those observed with SARS-CoV and MERS-CoV ^26-29^, with regard to lesion type and cell tropism.

Serological responses in humans are not typically detectable before 6 days after symptom onset, with IgG titers between 100 and 10,000 observed after 12 to 21 days^30,31^. Neutralizing titers were generally between 20 – 160. This corresponds to the results in our rhesus macaque model, where IgG responses were detected around 7-10 dpi and IgG titers peaked between 12 and 15 days at 1600 – 3200 with neutralizing titers between 10 and 60. This seroconversion was not directly followed by a decline in viral loads, as observed in COVID-19 patients^30,31^.

Taken together, we conclude that the rhesus macaque model recapitulates COVID-19, with clear evidence of virus replication in the upper and lower respiratory tract, shedding upper, lower respiratory and intestinal tract, the presence of pulmonary infiltrates, histological lesions resembling those observed with SARS-CoV and MERS-CoV and seroconversion. This extensive dataset allows us to bridge between the our Rhesus macaques model and the disease observed in humans and to utilize this animal model to assess the clinical and virological efficacy of medical countermeasures. We have therefore moved forward to test antiviral treatments and vaccines in this model.

## Methods

### Ethics and biosafety statement

All animal experiments were approved by the Institutional Animal Care and Use Committee of Rocky Mountain Laboratories, NIH and carried out by certified staff in an Association for Assessment and Accreditation of Laboratory Animal Care (AAALAC) International accredited facility, according to the institution’s guidelines for animal use, following the guidelines and basic principles in the NIH Guide for the Care and Use of Laboratory Animals, the Animal Welfare Act, United States Department of Agriculture and the United States Public Health Service Policy on Humane Care and Use of Laboratory Animals. Rhesus macaques were housed in adjacent individual primate cages allowing social interactions, in a climate-controlled room with a fixed light-dark cycle (12-hr light/12-hr dark). Animals were monitored at least twice daily throughout the experiment. Commercial monkey chow, treats, and fruit were provided twice daily by trained personnel. Water was available ad libitum. Environmental enrichment consisted of a variety of human interaction, manipulanda, commercial toys, videos, and music. The Institutional Biosafety Committee (IBC) approved work with infectious SARS-CoV-2 strains under BSL3 conditions. Sample inactivation was performed according to IBC-approved standard operating procedures for removal of specimens from high containment.

### Study design

To evaluate the use of rhesus macaques as a model for SARS-CoV-2, eight adult rhesus macaques (4 males, 4 females) were inoculated via a combination of intranasal (0.5ml per nostril), intratracheal (4ml), oral (1ml) and ocular (0.25ml per eye) of a 4×10^5^ TCID50/ml virus dilution in sterile DMEM. The animals were observed twice daily for clinical signs of disease using a standardized scoring sheet as described previously ^32^; the same person assessed the animals throughout the study. The predetermined endpoint for this experiment was 3 days post inoculation (dpi) for one group of 4 animals, and 21 dpi for the remaining 4 animals. Clinical exams were performed on 0, 1, 3, 5, 7, 10, 12, 14, 17 and 21 dpi on anaesthetized animals. On exam days, clinical parameters such as bodyweight, body temperature and respiration rate were collected, as well as ventro-dorsal and lateral chest radiographs. Chest radiographs were interpreted by a board-certified clinical veterinarian. The following samples were collected at all clinical exams: nasal, throat, urogenital and rectal swabs, blood. The total white blood cell count, lymphocyte, neutrophil, platelet, reticulocyte and red blood cell counts, hemoglobin, and hematocrit values were determined from EDTA blood with the IDEXX ProCyte DX analyzer (IDEXX Laboratories). Serum biochemistry (albumin, AST, ALT, GGT, BUN, creatinine) was analyzed using the Piccolo Xpress Chemistry Analyzer and Piccolo General Chemistry 13 Panel discs (Abaxis). During clinical exams on 1, 3, and 5 dpi bronchoalveolar lavages were performed using 10ml sterile saline. After euthanasia, necropsies were performed. The percentage of gross lung lesions was scored by a board-certified veterinary pathologist and samples of the following tissues were collected: inguinal lymph node, axillary lymph node, cervical lymph node, salivary gland, conjunctiva, nasal mucosa, oropharynx, tonsil, trachea, all six lung lobes, mediastinal lymph node, right and left bronchus, heart, liver, spleen, pancreas, adrenal gland, kidney, mesenteric lymph node, stomach, duodenum, jejunum, ileum, cecum, colon, urinary bladder, reproductive tract (testes or ovaries depending on sex of the animal), bone marrow, frontal brain, cerebellum and brainstem. Histopathological analysis of tissue slides was performed by a board-certified veterinary pathologist blinded to the group assignment of the animals.

### Virus and cells

SARS-CoV-2 isolate nCoV-WA1-2020 (MN985325.1)^18^ (Vero passage 3) was kindly provided by CDC and propagated once in VeroE6 cells in DMEM (Sigma) supplemented with 2% fetal bovine serum (Gibco), 1 mM L-glutamine (Gibco), 50 U/ml penicillin and 50 μg/ml streptomycin (Gibco) (virus isolation medium). The used virus stock was 100% identical to the initial deposited genbank sequence (MN985325.1) and no contaminants were detected. VeroE6 cells were maintained in DMEM supplemented with 10% fetal calf serum, 1 mM L-glutamine, 50 U/ml penicillin and 50 μg/ml streptomycin.

### Quantitative PCR

RNA was extracted from swabs and BAL using the QiaAmp Viral RNA kit (Qiagen) according to the manufacturer’s instructions. Tissues (30 mg) were homogenized in RLT buffer and RNA was extracted using the RNeasy kit (Qiagen) according to the manufacturer’s instructions. For detection of viral RNA, 5 µl RNA was used in a one-step real-time RT-PCR E assay^33^ using the Rotor-Gene probe kit (Qiagen) according to instructions of the manufacturer. In each run, standard dilutions of counted RNA standards were run in parallel, to calculate copy numbers in the samples. For detection of SARS-CoV-2 mRNA, primers targeting open reading frame 7 (ORF7) were designed as follows: forward primer 5’-TCCCAGGTAACAAACCAACC-3’, reverse primer 5’-GCTCACAAGTAGCGAGTGTTAT-3’, and probe FAM-ZEN-CTTGTAGATCTGTTCTCTAAACGAAC-IBFQ. 5 µl RNA was used in a one-step real-time RT-PCR using the Rotor-Gene probe kit (Qiagen) according to instructions of the manufacturer. In each run, standard dilutions of counted RNA standards were run in parallel, to calculate copy numbers in the samples.

### Histopathology and immunohistochemistry

Histopathology and immunohistochemistry were performed on rhesus macaque tissues. After fixation for a minimum of 7 days in 10 % neutral-buffered formalin and embedding in paraffin, tissue sections were stained with hematoxylin and eosin (HE). To detect SARS-CoV-2 antigen, immunohistochemistry was performed using an anti-SARS nucleocapsid protein antibody (Novus Biologicals) at a 1:250 dilution. Stained slides were analyzed by a board-certified veterinary pathologist.

#### Transmission electron microscopy

After fixation for 7 days with Karnovsky’s fixative at 4° C, excised tissues were post-fixed for 1 hour with 0.5% osmium tetroxide/0.8% potassium ferricyanide in 0.1 M sodium cacodylate, washed 3 × 5 minutes with 0.1M sodium cacodylate buffer, stained 1 hour with 1% tannic acid, washed with buffer and then further stained with2% osmium tetroxide in 0.1M sodium cacodylate and overnight with 1% uranyl acetate at 4° C. Specimens were dehydrated with a graded ethanol series with two final exchanges in 100% propylene oxide before infiltration and final embedding in Embed-812/Araldite resin. Thin sections were cut with a Leica EM UC6 ultramicrotome (Leica, Vienna, Austria), prior to viewing at 120 kV on a Tecnai BT Spirit transmission electron microscope (Thermo fisher/FEI, Hillsboro, OR). Digital images were acquired with a Gatan Rio bottom mount digital camera system (Gatan Inc., Pleasanton, CA and processed using Adobe Photoshop v. CC 2019 (Adobe Systems Inc, San Jose, CA).

#### Serum cytokine and chemokine analysis

Serum samples for analysis of cytokine/chemokine levels were inactivated with γ-radiation (2 MRad) according to standard operating procedures. Concentrations of granulocyte colony-stimulating factor, granulocyte-macrophage colony-stimulating factor, interferon (IFN)–γ, interleukin (IL)–1β, IL-1 receptor antagonist, IL-2, IL-4, IL-5, IL-6, IL-8, IL-10, IL-12/23 (p40), IL-13, IL-15, IL-17, MCP-1 and macrophage inflammatory protein (MIP)–1α, MIP-1β, soluble CD40-ligand (sCD40L), transforming growth factor-α, tumor necrosis factor (TNF)–α, vascular endothelial growth factor (VEGF) and IL-18 were measured on a Bio-Plex 200 instrument (Bio-Rad) using the Non-Human Primate Cytokine MILLIPLEX map 23-plex kit (Millipore) according to the manufacturer’s instructions.

### Serology

Sera were analyzed by SARS-CoV-2 spike protein (S) enzyme-linked immunosorbent assay (ELISA) as done previously for MERS-CoV^34^. Briefly, maxisorp (Nunc) plates were coated overnight with 100 ng/well S protein diluted in PBS^35^ (a kind gift of Barney Graham, Vaccine Research Center, NIH) and blocked with blocker casein in PBS (Life Technologies). Sera were serially diluted in duplicate. SARS-CoV-2-specific antibodies were detected using anti-monkey IgG polyclonal antibody HRP-conjugated antibody (KPL), peroxidase-substrate reagent (KPL) and stop reagent (KPL). Optical density (OD) was measured at 405⍰nm. The threshold of positivity was calculated by taking the average of the day 0 values multiplied by 3.

For neutralization, sera were heat-inactivated (30 min, 56 °C) and two-fold serial dilutions were prepared in 2% DMEM. Hereafter, 100 TCID_50_ of SARS-CoV-2 was added. After 60 min incubation at 37 °C, virus:serum mixture was added to VeroE6 cells and incubated at 37°C and 5% CO2. At 5 dpi, cytopathic effect was scored. The virus neutralization titer is expressed as the reciprocal value of the highest dilution of the serum which still inhibited virus replication. All sera were analyzed in duplicate.

### Data availability

All data are available on request.

## Acknowledgements

The authors would like to thank Susan Gerber and Natalie Thornburg (CDC) for providing the SARS-CoV-2 isolate used in this study; Barney Graham, Kizzmekia Corbett and Olubukola Abiona at the Vaccine Research Center (NIAID, NIH) for providing spike protein for serology; Anita Mora (NIAID, NIH) for help with figure design and staff of the Rocky Mountain Veterinary Branch (NIAID, NIH) for animal care. This study was supported by the Intramural Research Program, NIAID, NIH.

## Author contributions

VJM and EdW designed the study; VJM, FF,BW, NvD, LPP, JS, KMW, AO, JC, BB, VAA, RR, PH, GS, EF, DS and EdW acquired, analyzed and interpreted the data; VJM, PH, EF, DS and EdW wrote the manuscript. All authors have approved the submitted version.

## Competing interests

The authors declare no competing interests

