## Supplemental for "Respiratory disease and virus shedding in rhesus macaques inoculated with SARS-CoV-2"

**Table S1. Clinical signs observed in rhesus macaques inoculated with SARS-CoV-2.**

| **Animal** | **Clinical signs observed 1-6 dpi** | **Clinical signs observed 7-21 dpi** | **Observations at necropsy*** |
| --- | --- | --- | --- |
| **RM1** | Hunched posture; piloerection; tachypnea; flushed appearance; red eyes; very agitated; reduced appetite; mildly dehydrated.  Euthanized 3 dpi. | N/A | Gross lung lesions.  Enlarged tonsils and mediastinal lymph nodes.  Fluid-filled stomach, small and large intestine. |
| **RM2** | Piloerection; dyspnea; reduced appetite.  Euthanized 3 dpi. | N/A | Fluid-filled stomach, small and large intestine. |
| **RM3** | Piloerection; tachypnea; flushed appearance; reduced appetite; mildly dehydrated.  Euthanized 3 dpi. | N/A | Epistaxis. Gross lung lesions.  Enlarged mediastinal lymph nodes.  Fluid-filled stomach, small and large intestine. |
| **RM4** | Hunched posture; piloerection; tachypnea; dyspnea; reduced appetite.  Euthanized 3 dpi. | N/A | Gross lung lesions. Foamy exudate from trachea.  Enlarged mediastinal lymph nodes.  Fluid-filled stomach, small and large intestine. |
| **RM5** | Hunched posture; piloerection; tachypnea; dyspnea; reduced appetite. | Tachypnea; dyspnea; reduced appetite; mildly dehydrated.  Recovered on 9 dpi. | Gross lung lesions.  Enlarged mesenteric lymph nodes. |
| **RM6** | Hunched posture; piloerection; tachypnea; dyspnea; reduced appetite. | Piloerection; bradypnea; mildly dehydrated.  Recovered on 10 dpi. | None. |
| **RM7** | Hunched posture; piloerection; pale appearance; tachypnea; dyspnea; irregular; labored respirations; anorexia; mildly dehydrated. | Hunched posture; piloerection; pale appearance; tachypnea; dyspnea; reduced appetite; mildly dehydrated.  Recovered on 17 dpi. | None. |
| **RM8** | Hunched posture; piloerection; pale appearance; increased, dyspnea; reduced appetite. | Hunched posture; piloerection; pale appearance; increased, dyspnea; nasal discharge; reduced appetite; mildly dehydrated.  Recovered on 13 dpi. | Gross lung lesions. |

* Incidental observations not related to coronavirus infection were omitted from this table.

**Table S2. Blood chemistry in rhesus macaques infected with SARS-CoV-2.** Blood samples were analyzed using the General Chemistry 13 panel in a Piccolo Express chemistry analyzer.

| **Animal** | **Time (dpi)** | **Glucose (mg/dL)** | **BUN (mg/dL)** | **Creatinine (mg/dL)** | **Calcium (mg/dL)** | **Albumin (g/dL)** | **Total protein (g/dL)** | **Calc Glob (g/dL)** | **ALT (U/L)** | **AST (U/L)** | **AST/ALT Ratio** | **ALP (U/L)** | **Total bilirubin (mg/dL)** | **GGT (U/L)** | **Amylase (U/L)** | **Hem** | **Lip** | **Ict** |
| --- | --- | --- | --- | --- | --- | --- | --- | --- | --- | --- | --- | --- | --- | --- | --- | --- | --- | --- |
| **RM1** | **0** | 118 | 15 | 1.1 | 9.5 | 3.2 | 6.6 | 3.4 | 46 | 47 | 1.02 | 516 | 0.6 | 102 | 442 | 0 | 0 | 0 |
|  | **1** | 124 | 14 | 1.1 | 9.1 | 3.1 | 6.2 | 3.1 | 83 | 66 | 0.80 | 463 | 0.7 | 89 | 591 | 1 | 0 | 0 |
|  | **3** | 93 | 20 | 1.1 | 8.6 | 3.0 | 6.1 | 3.1 | 78 | 57 | 0.73 | 411 | 0.7 | 81 | 464 | 0 | 0 | 0 |
| **RM2** | **0** | 97 | 11 | 0.6 | 10.5 | 3.4 | 7.5 | 4.1 | 31 | 29 | 0.94 | 161 | 0.6 | 53 | 325 | 0 | 0 | 0 |
|  | **1** | 132 | 10 | 0.8 | 10.6 | 3.3 | 7.5 | 4.2 | 29 | 29 | 1.00 | 161 | 0.5 | 50 | 373 | 2 | 0 | 0 |
|  | **3** | 85 | 8 | 0.7 | 10.0 | 3.3 | 7.4 | 4.1 | 25 | 33 | 1.32 | 172 | 0.7 | 49 | 377 | 1 | 0 | 0 |
| **RM3** | **0** | 54 | 22 | 0.9 | 10.1 | 3.5 | 6.7 | 3.2 | 36 | 44 | 1.22 | 767 | 0.6 | 127 | 330 | 0 | 0 | 0 |
|  | **1** | 104 | 21 | 0.9 | 9.6 | 3.5 | 6.5 | 3.0 | 46 | 66 | 1.43 | 701 | 0.8 | 117 | 539 | 0 | 0 | 0 |
|  | **3** | 82 | 20 | 1.1 | 9.4 | 3.5 | 6.6 | 3.1 | 44 | 41 | 0.93 | 537 | 0.7 | 106 | 324 | 0 | 0 | 0 |
| **RM4** | **0** | 84 | 19 | 0.9 | 10.1 | 3.5 | 7.2 | 3.7 | 70 | 49 | 0.70 | 184 | 0.5 | 75 | 288 | 0 | 0 | 0 |
|  | **1** | 85 | 15 | 0.6 | 9.8 | 3.2 | 6.5 | 3.3 | 76 | 67 | 0.88 | 166 | 0.5 | 65 | 251 | 0 | 1 | 0 |
|  | **3** | 66 | 19 | 0.9 | 8.7 | 3.3 | 6.5 | 3.2 | 58 | 39 | 0.67 | 158 | 0.5 | 60 | 249 | 0 | 0 | 0 |
| **RM5** | **0** | 77 | 12 | 0.8 | 9.8 | 3.3 | 6.6 | 3.3 | 37 | 38 | 1.03 | 546 | 0.5 | 98 | 345 | 0 | 0 | 0 |
|  | **1** | 103 | 14 | 0.9 | 10.6 | 3.5 | 7.7 | 4.2 | 56 | 52 | 0.93 | 598 | 0.6 | 106 | 385 | 1 | 0 | 0 |
|  | **3** | 78 | 15 | 1.0 | 9.3 | 3.1 | 6.9 | 3.8 | 40 | 48 | 1.20 | 457 | 0.6 | 87 | 317 | 0 | 0 | 0 |
|  | **5** | 85 | 14 | 0.9 | 10.3 | 3.4 | 7.8 | 4.4 | 80 | 62 | 0.78 | 459 | 0.6 | 95 | 327 | 0 | 0 | 0 |
|  | **7** | 75 | 15 | 0.7 | 9.5 | 3.1 | 6.7 | 3.6 | 55 | 39 | 0.71 | 415 | 0.6 | 84 | 341 | 0 | 0 | 0 |
|  | **10** | 75 | 18 | 0.8 | 9.7 | 3.1 | 7.0 | 3.9 | 42 | 35 | 0.83 | 461 | 0.5 | 84 | 352 | 0 | 0 | 0 |
|  | **12** | 85 | 13 | 0.9 | 9.6 | 3.2 | 7.3 | 4.1 | 48 | 38 | 0.79 | 505 | 0.6 | 87 | 370 | 0 | 0 | 0 |
|  | **14** | 79 | 15 | 0.8 | 9.5 | 3.1 | 6.7 | 3.6 | 51 | 35 | 0.69 | 446 | 0.5 | 80 | 408 | 0 | 0 | 0 |
|  | **17** | 82 | 16 | 0.9 | 10.0 | 3.3 | 6.8 | 3.5 | 83 | 45 | 0.54 | 464 | 0.5 | 84 | 383 | 0 | 0 | 0 |
|  | **21** | 73 | 8 | 0.8 | 9.2 | 3.2 | 6.3 | 3.1 | 60 | 50 | 0.83 | 450 | 0.4 | 81 | 333 | 0 | 0 | 0 |
| **RM6** | **0** | 81 | 15 | 0.7 | 9.6 | 3.3 | 5.9 | 2.6 | 28 | 29 | 1.04 | 361 | 0.5 | 79 | 354 | 0 | 0 | 0 |
|  | **1** | 118 | 18 | 1.0 | 11.8 | 4.6 | 8.4 | 3.8 | 41 | 45 | 1.10 | 442 | 0.6 | 101 | 698 | 0 | 0 | 0 |
|  | **3** | 81 | 12 | 0.7 | 9.8 | 3.5 | 6.8 | 3.3 | 35 | 32 | 0.91 | 302 | 0.5 | 80 | 383 | 0 | 0 | 0 |
|  | **5** | 86 | 15 | 0.8 | 11.0 | 3.9 | 8.0 | 4.1 | 42 | 35 | 0.83 | 331 | 0.5 | 94 | 382 | 0 | 0 | 0 |
|  | **7** | 68 | 15 | 0.8 | 10.3 | 3.7 | 7.1 | 3.4 | 42 | 32 | 0.76 | 313 | 0.5 | 86 | 330 | 0 | 0 | 0 |
|  | **10** | 72 | 16 | 0.7 | 10.2 | 3.7 | 7.2 | 3.5 | 50 | 29 | 0.58 | 320 | 0.5 | 90 | 399 | 0 | 0 | 0 |
|  | **12** | 76 | 13 | 0.7 | 10.4 | 3.7 | 7.2 | 3.5 | 44 | 28 | 0.64 | 326 | 0.5 | 90 | 382 | 0 | 0 | 0 |
|  | **14** | 72 | 14 | 0.7 | 10.5 | 3.8 | 7.3 | 3.5 | 41 | 30 | 0.73 | 323 | 0.5 | 90 | 411 | 0 | 0 | 0 |
|  | **17** | 74 | 17 | 0.7 | 10.7 | 3.8 | 7.5 | 3.7 | 37 | 31 | 0.84 | 305 | 0.5 | 87 | 389 | 0 | 0 | 0 |
|  | **21** | 68 | 13 | 0.7 | 10.3 | 3.7 | 7.0 | 3.3 | 49 | 38 | 0.78 | 316 | 0.4 | 84 | 353 | 0 | 0 | 0 |
| **RM7** | **0** | 98 | 11 | 0.9 | 10.1 | 3.8 | 7.3 | 3.5 | 45 | 33 | 0.73 | 770 | 0.5 | 136 | 417 | 0 | 0 | 0 |
|  | **1** | 143 | 14 | 1.0 | 10.9 | 4.0 | 7.9 | 3.9 | 60 | 58 | 0.97 | 716 | 0.6 | 135 | 342 | 0 | 0 | 0 |
|  | **3** | 90 | 13 | 0.8 | 9.2 | 3.3 | 6.6 | 3.3 | 44 | 37 | 0.84 | 468 | 0.5 | 103 | 384 | 0 | 0 | 0 |
|  | **5** | 102 | 17 | 0.9 | 11.4 | 3.9 | 8.2 | 4.3 | 46 | 37 | 0.80 | 501 | 0.5 | 118 | 489 | 0 | 0 | 0 |
|  | **7** | 69 | 13 | 0.8 | 9.4 | 3.3 | 6.9 | 3.6 | 42 | 29 | 0.69 | 424 | 0.5 | 95 | 421 | 0 | 0 | 0 |
|  | **10** | 75 | 17 | 0.7 | 10.0 | 3.5 | 7.1 | 3.6 | 43 | 28 | 0.65 | 478 | 0.5 | 99 | 448 | 0 | 0 | 0 |
|  | **12** | 83 | 14 | 1.0 | 10.7 | 4.0 | 8.1 | 4.1 | 46 | 38 | 0.83 | 604 | 0.5 | 116 | 457 | 0 | 0 | 0 |
|  | **14** | 72 | 12 | 0.8 | 9.8 | 3.4 | 6.9 | 3.5 | 84 | 33 | 0.39 | 540 | 0.5 | 100 | 481 | 0 | 0 | 0 |
|  | **17** | 77 | 17 | 1.0 | 10.1 | 3.5 | 6.9 | 3.4 | 60 | 53 | 0.88 | 513 | 0.5 | 99 | 441 | 0 | 0 | 0 |
|  | **21** | 67 | 9 | 0.7 | 9.7 | 3.5 | 6.7 | 3.2 | 46 | 41 | 0.89 | 569 | 0.4 | 105 | 416 | 0 | 0 | 0 |
| **RM8** | **0** | 110 | 14 | 0.8 | 11.2 | 3.6 | 7.6 | 4.0 | 26 | 31 | 1.19 | 260 | 0.5 | 79 | 444 | 0 | 0 | 0 |
|  | **1** | 110 | 11 | 0.8 | 11.0 | 3.6 | 7.4 | 3.8 | 46 | 57 | 1.24 | 259 | 0.6 | 75 | 493 | 0 | 0 | 0 |
|  | **3** | 76 | 12 | 0.7 | 8.9 | 3.2 | 6.7 | 3.5 | 58 | 49 | 0.84 | 197 | 0.6 | 64 | 438 | 0 | 0 | 0 |
|  | **5** | 83 | 13 | 0.7 | 11.1 | 3.7 | 7.8 | 4.1 | 68 | 36 | 0.53 | 202 | 0.5 | 73 | 467 | 1 | 0 | 0 |
|  | **7** | 75 | 13 | 0.7 | 9.6 | 3.3 | 6.7 | 3.4 | 48 | 31 | 0.65 | 182 | 0.5 | 62 | 398 | 0 | 0 | 0 |
|  | **10** | 76 | 14 | 0.6 | 10.1 | 3.5 | 6.7 | 3.2 | 38 | 29 | 0.76 | 193 | 0.5 | 61 | 383 | 2 | 0 | 0 |
|  | **12** | 76 | 11 | 0.7 | 9.2 | 3.4 | 6.7 | 3.3 | 34 | 27 | 0.79 | 211 | 0.5 | 61 | 359 | 0 | 0 | 0 |
|  | **14** | 70 | 11 | 0.7 | 9.7 | 3.5 | 6.7 | 3.2 | 34 | 28 | 0.82 | 229 | 0.5 | 65 | 447 | 0 | 0 | 0 |
|  | **17** | 77 | 12 | 0.7 | 10.1 | 3.4 | 6.8 | 3.4 | 34 | 36 | 1.06 | 206 | 0.5 | 62 | 436 | 0 | 0 | 0 |
|  | **21** | 74 | 11 | 0.7 | 9.9 | 3.6 | 6.6 | 3.0 | 35 | 27 | 0.77 | 207 | 0.5 | 62 | 367 | 0 | 0 | 0 |

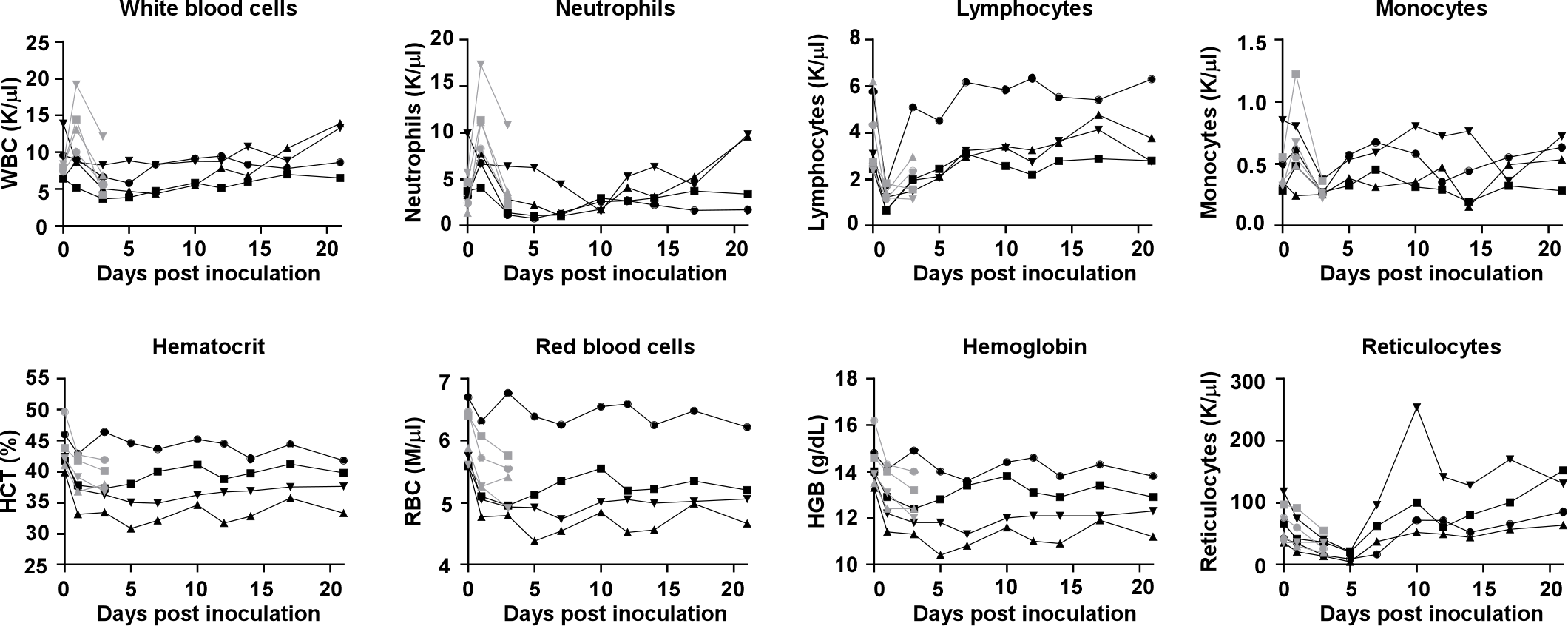

**Figure S1. Hematological changes in rhesus macaques infected with SARS-CoV-2.** Identical symbols have been used to denote identical animals throughout the figures in this manuscript.

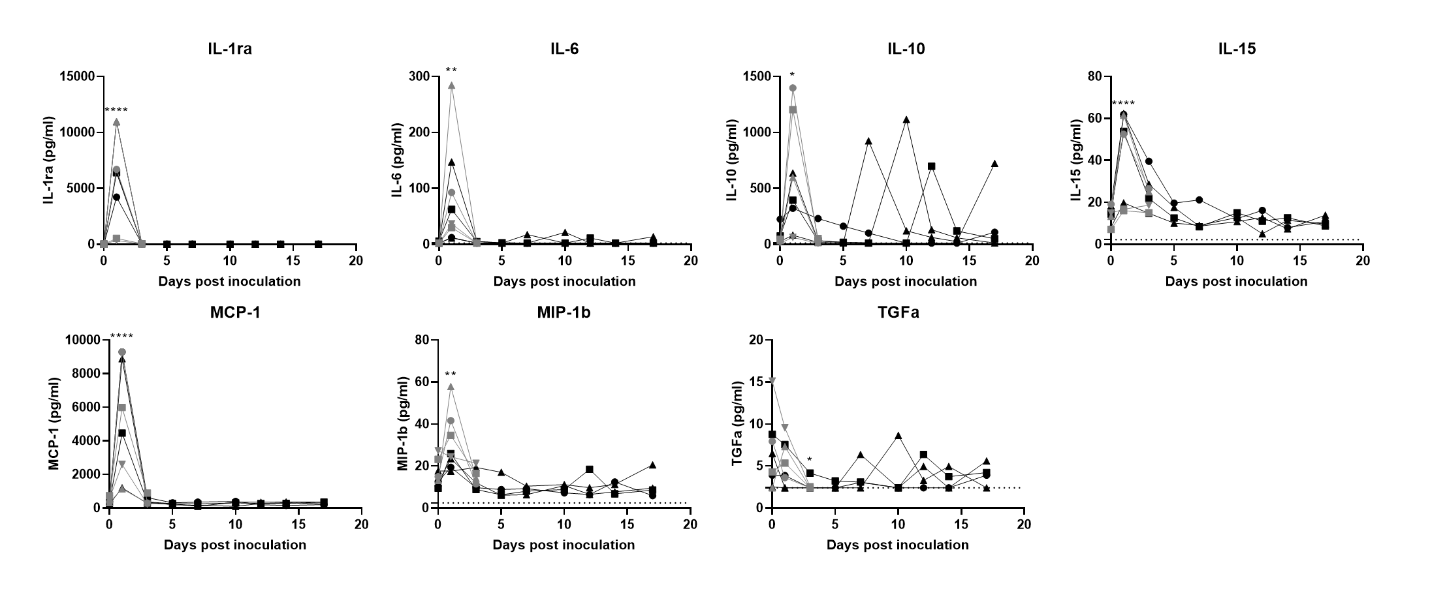

**Figure S2. Cytokine and chemokine levels in serum of rhesus macaques infected with SARS-CoV-2.** The levels of 23 cytokines and chemokines were determined in serum at different timepoints after inoculation. Levels are displayed only for those cytokines and chemokines where statistically significant (1-way ANOVA with multiple comparisons) were observed compared to levels on day of inoculation. Identical symbols have been used to denote identical animals throughout the figures in this manuscript. The lower limit of detection is indicated with a dotted line. *P<0.05; ** P<0.01; ****P<0.0001.

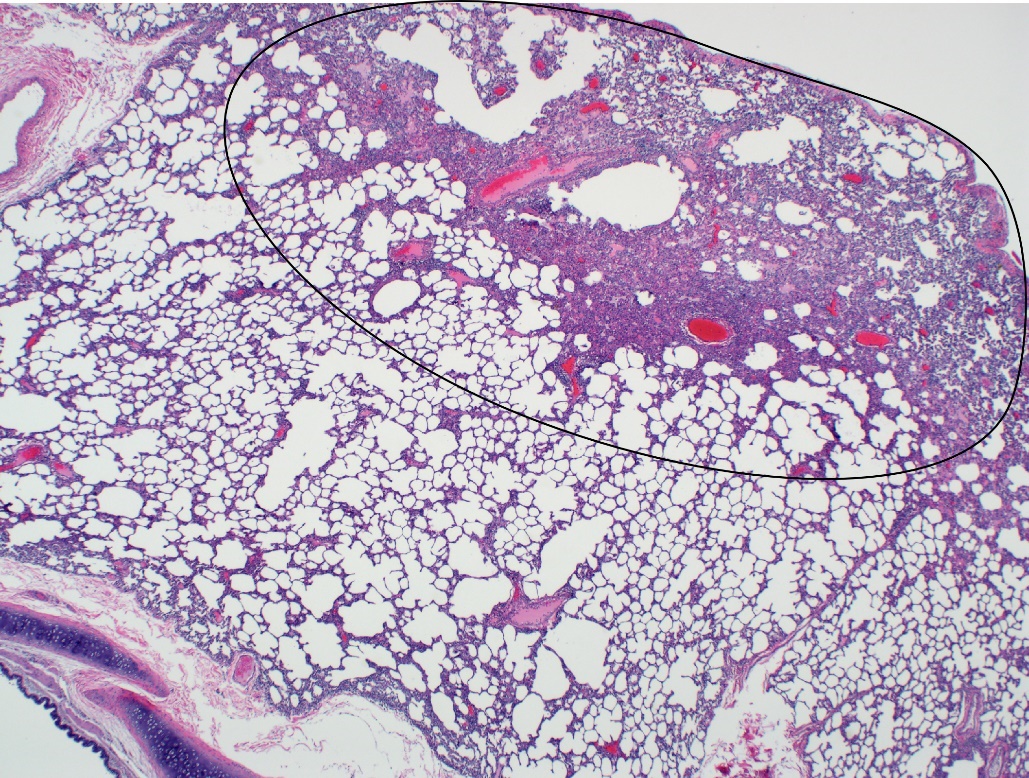

**Figure S3. Focal lesion in lung of a rhesus macaque infected with SARS-CoV-2.** This low magnification figure displays the focal nature of SARS-CoV-2 lesions in the lungs of animals euthanized on 3 dpi. The circle indicates the lung affected by lesion; the remaining lung tissue is healthy.

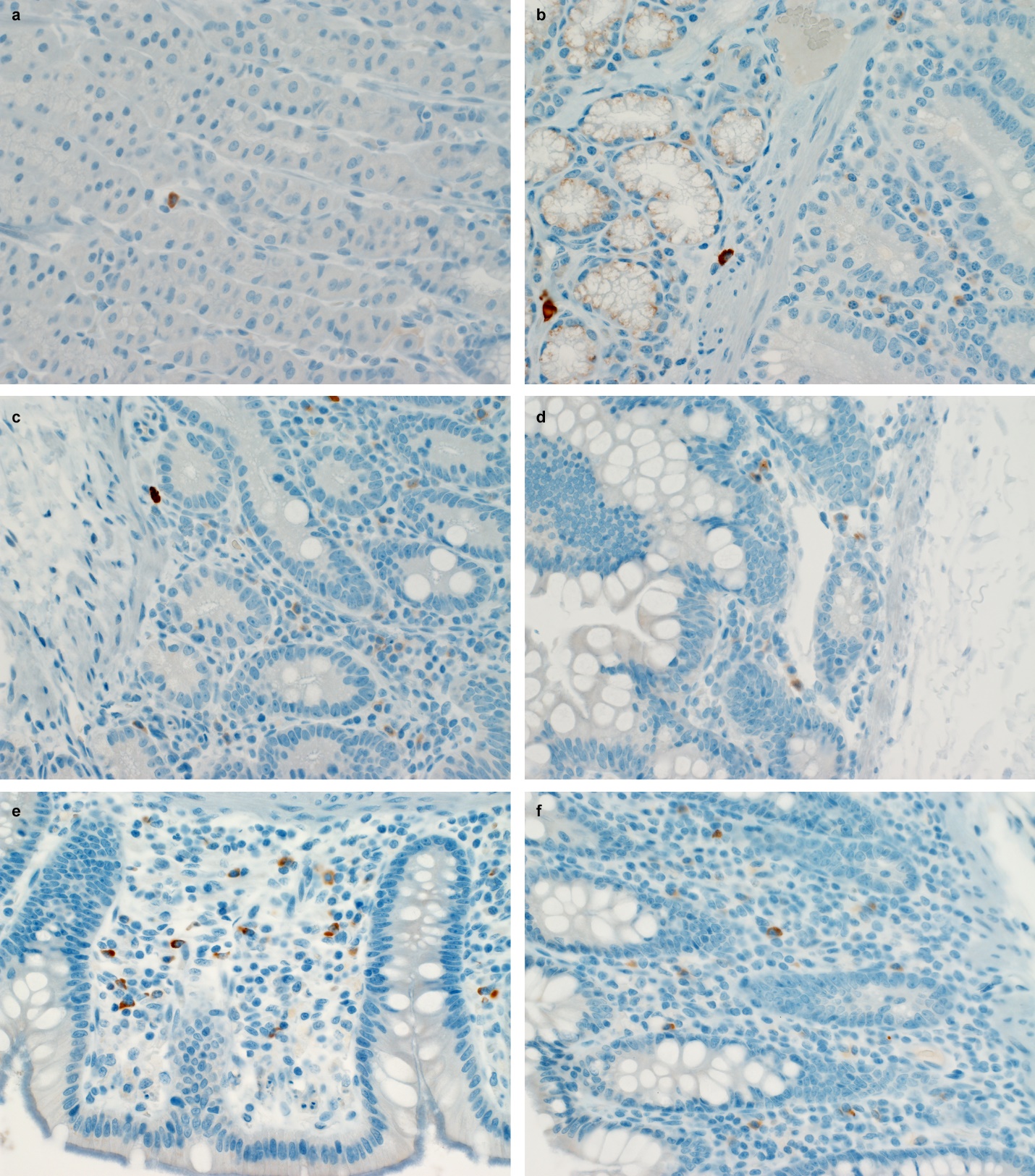

**Figure S4. SARS-CoV-2 antigen in the gastrointestinal tract of a rhesus macaque infected with SARS-CoV-2.** Mononuclear cells staining positive for SARS-CoV-2 antigen in the lamina propria of stomach (a), duodenum (b), jejunum (c), ileum (d), cecum (e) and colon (f) of an animal infected with SARS-CoV-2 and euthanized on 3 dpi.

**Figure S5. Viral loads detected in tissues collected on 21 dpi from rhesus macaques infected**
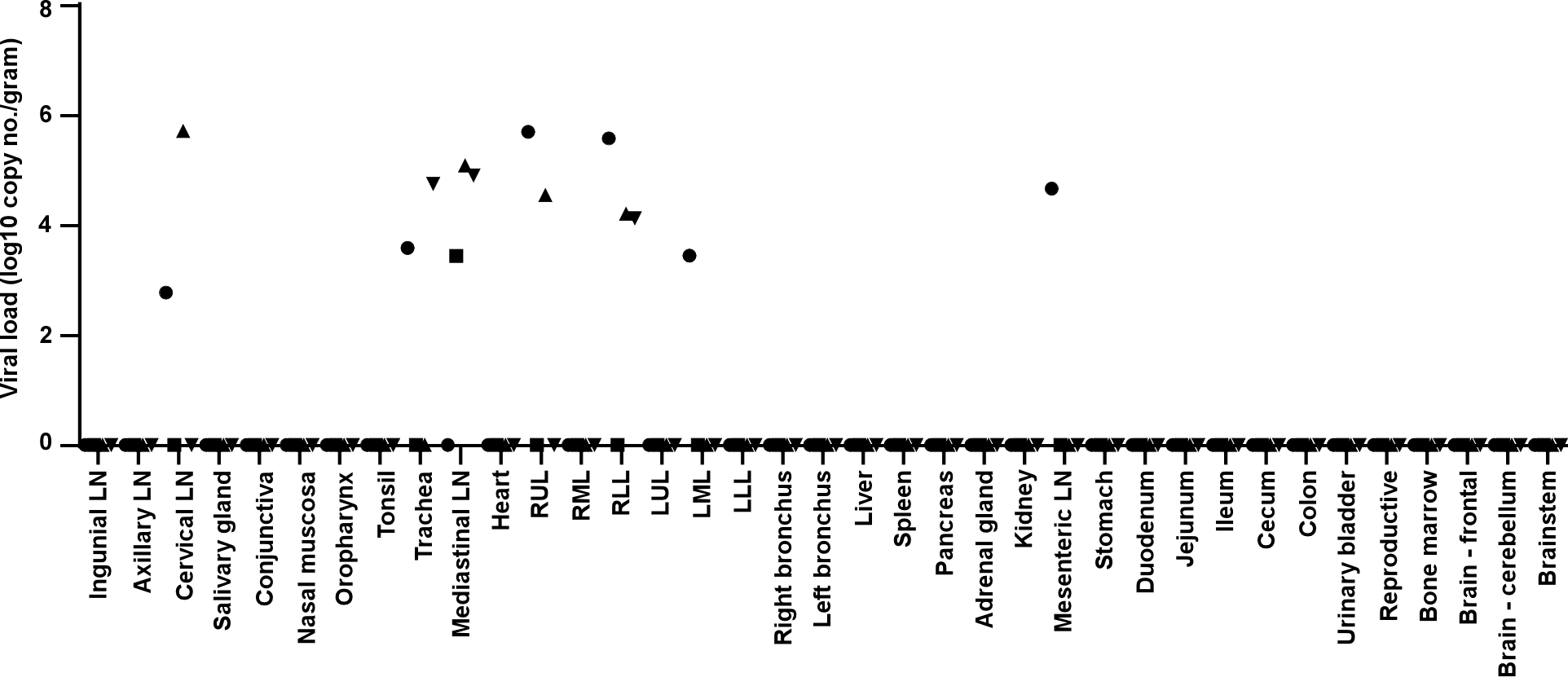
**with SARS-CoV-2.** Four adult rhesus macaques were inoculated with SARS-CoV-2 isolate nCoV-WA1-2020 and euthanized on 3 dpi. Thirty-seven tissues were collected at necropsy and analyzed for the presence of viral RNA by qRT-PCR. Identical symbols have been used to denote identical animals throughout the figures in this manuscript. LN: lymph node; RUL: right upper lung lobe; RML: right middle lung lobe; RLL: right lower lung lobe; LUL: left upper lung lobe; LML: left middle lung lobe; LLL: left lower lung lobe.

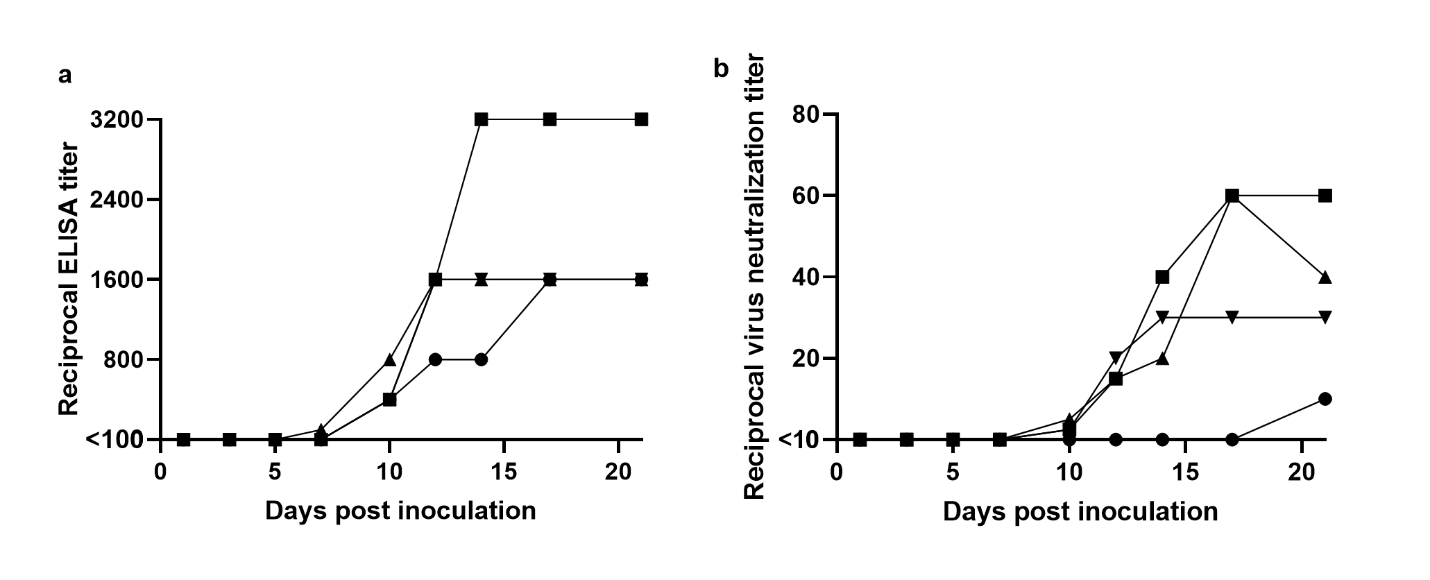

**Figure S6. Antibody response in rhesus macaques infected with SARS-CoV-2.** Sera collected after inoculation were tested for the presence of IgG against SARS-CoV-2 spike in ELISA (a) and for the presence of neutralizing antibodies ina microneutralization assay (b). All sera were analyzed in duplicate. Identical symbols have been used to denote identical animals throughout the figures in this manuscript.
